# Cognitive and white-matter compartment models revealed the contribution of microstructural variability along sensorimotor tracts to simple reaction time

**DOI:** 10.1101/473660

**Authors:** Esin Karahan, Alison G. Costigan, Kim S. Graham, Andrew D. Lawrence, Jiaxiang Zhang

**Author notes:** Correspondence should be addressed to: Jiaxiang Zhang, Cardiff University Brain Research Imaging Centre, Maindy Road, Cardiff CF24 4HQ, UK.

## Abstract

The speed of voluntary reaction to an external stimulus varies substantially between individuals and is impaired in ageing. However, the neuroanatomical origins of inter-individual variability in reaction time (RT) remain largely unknown. Here, we combined a cognitive model of RT and a biophysical compartmental model of diffusion-weighted MRI (DWI) to characterize the relationship between RT and microstructure of the corticospinal tract (CST) and the optic radiation (OR), the primary motor output and visual input pathways associated with visual-motor responses.

We fitted an accumulator model of RT to 46 female participants’ behavioral performance in a simple reaction time task. The non-decision time parameter (*T_er_*) derived from the model was used to account for the latencies of stimulus encoding and action initiation. From multi-shell DWI data, we quantified tissue microstructure of the CST and OR with the neurite orientation dispersion and density imaging (NODDI) model as well as the conventional diffusion tensor imaging (DTI) model.

Using novel skeletonization and segmentation approaches, we showed that DWI-based microstructure metrics varied substantially along CST and OR. The *T_er_* of individual participants was negatively correlated with the NODDI measure of the neurite density in the bilateral superior CST. At an uncorrected threshold, the *T_er_* positively correlated with the DTI measure of fractional anisotropy in an anterior segment of left OR. Further, we found no significant correlation between the microstructural measures and mean RT. Thus, our findings suggest a link between the inter-individual variability of sensorimotor speed and selective microstructural properties in white matter tracts.

## Introduction

Voluntary response to external stimuli is a hallmark of cognitive control that encompasses perceptual, decision and motor processes. The reaction time (RT), measured as the latency between a preparatory stimulus and a pre-defined action, varies substantially across individuals (Jensen, 2006), changes during development (Dykiert et al., 2012), ageing (Woods et al., 2015) and neurodegeneration (Gorus et al., 2008), and have implications for mortality (Der and Deary, 2018). RT has also been identified as a marker of mental processing speed (Ho et al., 1988; Sheppard and Vernon, 2008), a heritable trait relating to intelligence (Vernon, 1989; Rabbitt, 1996).

Individual differences in RT have been regarded as reflections of a “primitive” neurophysiological characteristic (Rabbitt et al., 2001), and the potential of RT as a behavioral phenotype requires to understand its microstructural underpinnings (Turken et al., 2008; Penke et al., 2010). In humans, the diffusion tensor imaging (DTI) is commonly used to estimate tissue microstructure from diffusion-weighted MRI (DWI) (Basser and Pierpaoli, 1996), which is sensitive to the degree of anisotropic water diffusion due to cellular structures. The DTI measures of several white matter pathways have been shown to correlate with RT, both in adults (Tuch et al., 2005; Turken et al., 2008; Konrad et al., 2009; Mayer and Vuong, 2014) and children (Madsen et al., 2011; Tamnes et al., 2012; Scantlebury et al., 2014), possibly owing to the difference in experiments and cohorts across studies. One hypothesis of these structural-functional correlations is that the inter-individual variability in RT is due to variations in tissue microstructure, such as axon diameter or myelination that affect the nerve conduction velocity (Waxman, 1980; Fields, 2008; Seidl, 2014; Buskey et al., 2017) and in turn affect RT (Chevalier et al., 2015; Chopra et al., 2018).

However, research on the relationship between RT and DTI raises two unresolved issues. First, RT, even in simple behavioral tasks, is a hybrid measure of multiple intermixed cognitive processes (Gold and Shadlen, 2007; Forstmann et al., 2016; Ratcliff et al., 2016). Second, the conventional DTI model cannot distinguish microstructural properties between intra-and extra-cellular space, because it assumes a single tissue compartment (Beaulieu, 2002).

This study addressed these questions by combining a cognitive model of RT and a biophysical compartment model of the DWI signal. We used the cognitive model (Brown and Heathcote, 2008) to decompose a non-decision time measure from individual participant’s RT distribution in a simple reaction time task, which accounts for the latencies of stimulus encoding and action initiation (Lo and Wang, 2006; Donkin et al., 2011). DWI data was analyzed with the Neurite Orientation Dispersion and Density Imaging (NODDI) model (Zhang et al., 2012a). NODDI estimates separately the isotropic and anisotropic diffusion in multiple compartments, allowing two specific measures of tissue microstructure: the neurite density index (NDI) as the intra-cellular volume fraction and the orientation dispersion index (ODI) explaining the bending or fanning of axon orientations.

Furthermore, using probabilistic tractography and volume skeletonization, we developed a new method to quantify the changes of microstructural metrics along white matter tracts. We focused on two *a priori* tracts according to the functional anatomy of voluntary movement and visual processing that are relevant to the simple reaction time task. The first tract was the corticospinal tract (CST), a major output pathway carrying motor impulses from the giant pyramidal cells in the motor area to the midbrain, vital for voluntary movements (Bortoff and Strick, 1993; Lemon, 2008). The second was the optic radiation (OR), the prominent white matter relay in the visual system, transmitting information from the lateral geniculate nucleus to V1 (Garey and Powell, 1971; Ebeling and Reulen, 1988).

Our results demonstrated that microstructural metrics varied substantially along CST and OR. Higher neurite density in bilateral superior CST was associated with faster non-decision time across participants, but not with the mean RT. These findings suggested the existence of microstructural-specific influence of inter-individual variability in subcomponents of the action decision process.

## Materials and Methods

### Participants

Forty-six healthy young adults participated in the study (all females, age range 19-24 years; mean age 20.8 years) following written informed consent. Participants had normal or corrected-to-normal vision, and none reported a history of neurological or psychiatric illness. This study was approved by the Cardiff University School of Psychology Research Ethics Committee.

### Experimental design

Participants performed a visually paced simple reaction time (SRT) task adapted from previous studies (Zhang et al., 2012b; Shafto et al., 2014). The task was conducted in a behavioral testing room. The participants were presented with an image of a right hand on a 24-inch LED monitor with 1920×1080 screen resolution (ASUS VG248QE) and pressed the spacebar on the keyboard button with their right index finger. Four transparent circles superimposed above the four fingers in the hand image to serve as task cues (Fig. 1A). On each trial, the task cue above the index finger in the image turned to an opaque circle, indicating the start of the trial, and the participants were instructed to respond as quickly as possible. After participants’ response or after a maximum of 3 seconds response window, the opaque task cue extinguished and changed back to a transparent circle. Visual stimuli were presented by using Visual Basic 5.0.

**Figure 1.**
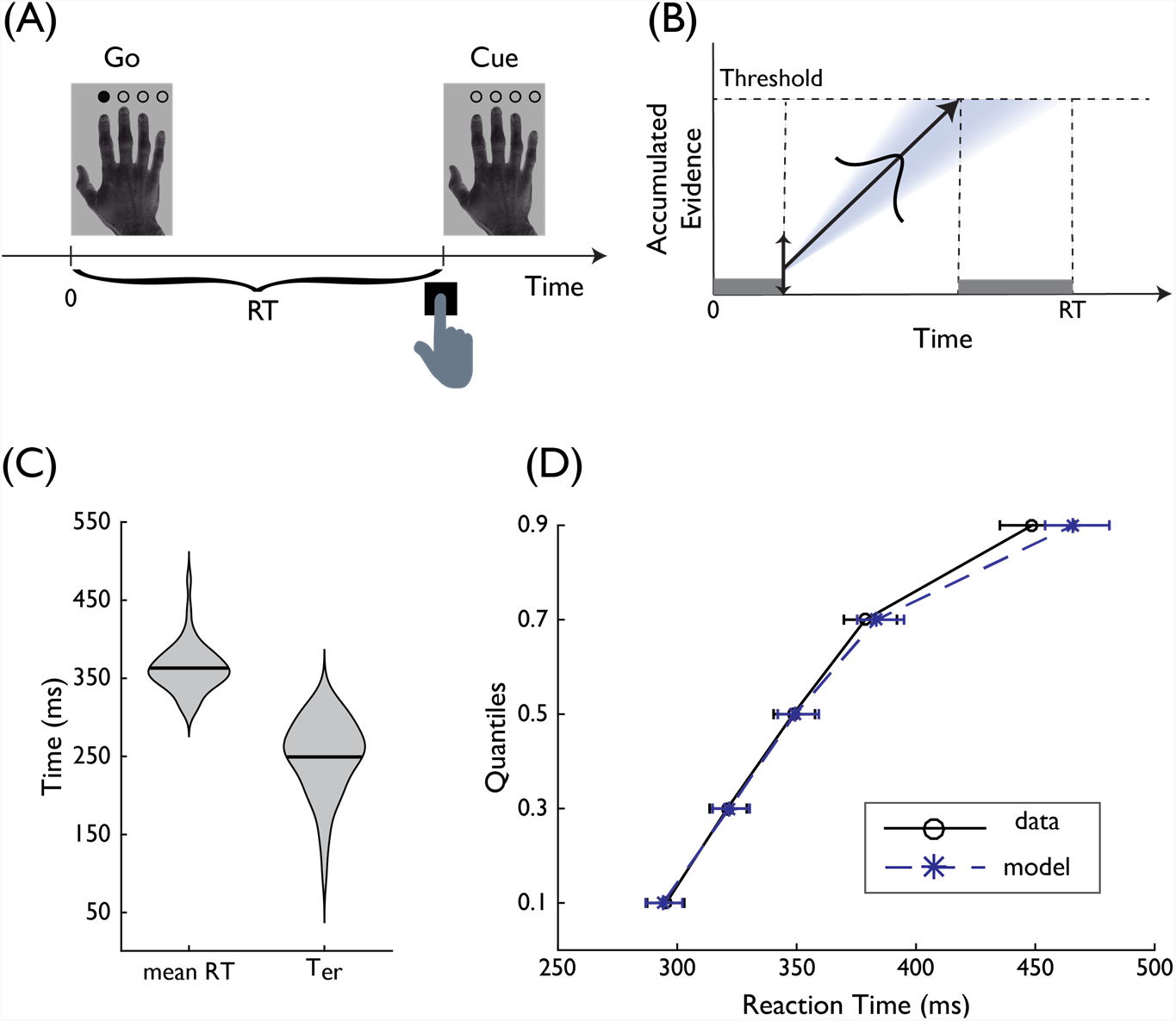
**(A)** Experimental paradigm of the simple reaction time (SRT) task. Participants were instructed to respond when a filled circle appeared over the index finger in the hand picture. **(B)** Exemplar time course of the linear ballistic accumulator (LBA) model. The LBA assumes that accumulated evidence for an action decision is accumulated linearly over time, and a decision is made once the accumulated evidence reaches a threshold. In each trial, the starting point of the accumulation process is sampled from a uniform distribution. The rate of accumulation is sampled from a Gaussian. The model predicted RT includes the duration of the accumulation process and a non-decision time (*T_er_*), which accounts for the latencies of non-decision processes such as stimulus encoding and action initiation which are shown as shaded area. **(C)** Violin plots (mean and density) of the mean RT and *T_er_* across participants. **(D)** The fit of the LBA model to RT quantiles in the SRT task. Error bars denote the 95% confidence intervals.

After a short practice, the participants performed 50 trials of the SRT task. To discourage proactive response strategies, the inter-trial interval randomly varied across trials with a skewed distribution (minimum 1800 ms, maximum 6800 ms, and mean 3700 ms). Reaction time (RT) was measured as the latency between task cue onset and button press. Trials with RT shorter than 150 ms or larger than 1500 ms were excluded from further analysis (0.65 % of the total number of trials across all participants). Mean RT was then calculated as the behavioral dependent measure for each participant.

### Accumulator model of simple actions and parameter estimation

We further analyzed the RT data using a cognitive model of RT, the linear ballistic accumulator (LBA) model (Brown and Heathcote, 2008). The LBA model is a simplified implementation of a large family of sequential sampling models (Ratcliff and Smith, 2004; Bogacz et al., 2006; Gold and Shadlen, 2007; Zhang, 2012) and has been used to study the cognitive processes underlying decision making (Ho et al., 2009; Forstmann et al., 2011) and action selection (Zhang et al., 2012b).

For the SRT task with one possible action, we assumed that the process is governed by a linear evidence accumulation process (Fig. 1B), from a randomly sampled starting point to a decision threshold *B* (see also (Ratcliff and Van Dongen, 2011) and (Schurger et al., 2012) for similar approaches). The speed of evidence accumulation varies across trials as a Gaussian random variable with a mean *µ* and standard deviation *σ*. The model-predicted RT is given by the duration of the accumulation process to reach a decision threshold *B*, plus a constant non-decision time *T_er_*. The non-decision time *T_er_* does not relate to evidence accumulation, but accounts for the latency of other processes including motor response initiation and stimulus encoding (Gold and Shadlen, 2007; Brown and Heathcote, 2008).

We fitted the LBA model to the RT distribution from the SRT task using a minimization procedure validated in previous studies of RT modelling (Bogacz and Cohen, 2004; Bogacz et al., 2006; Boucher et al., 2007; Dean et al., 2011; Zhang et al., 2012b, 2016). For each participant, the observed RT distribution was binned into the 0.1, 0.3, 0.5, 0.7, and 0.9 quantiles (Ratcliff and Smith, 2004), and the model prediction of the five RT quantiles were estimated from 100,000 numerical simulations. The starting point variability was fixed at 0.5 as the scaling parameter (Brown and Heathcote, 2008; Donkin et al., 2009). The model parameters (*T_er_*, *B*, *µ*, and *σ*) were determined by minimizing the likelihood ratio chi-square statistic between the observed and predicted RT distributions using the Simplex search algorithm (Nelder and Mead, 1965). To optimize the chance of locating the optimal model parameters, the minimization procedure started with a set of initial parameter values. Each initial parameter set was chosen from 100 randomly generated values that produced the best fit. The entire minimization procedure was then repeated 20 iterations to identify the best-fitting model parameters. The two time-dependent measures, the non-decision time *T_er_* and the mean RT, were then associated with microstructural metrics from diffusion MRI.

### MRI data acquisition

Whole-brain two-shell DWI data were acquired using a Siemens 3T Prisma MRI scanner and a 32-channel receiver head coil (Siemens Medical Systems, Germany) at the Cardiff University Brain Research Imaging Centre (CUBRIC) with single-shot spin-echo echo-planar imaging pulse sequence (echo time 67 ms, repetition time 9400 ms, field of view 256×256 mm, acquisition matrix size 128×128, voxel size 2×2×2 mm). Diffusion sensitizing gradients were applied in 30 isotropic directions at a *b*-value of 1200 s/mm^2^ and in 60 isotropic directions at a *b*-value of 2400 s/mm^2^. Six images with no diffusion weighting (*b* = 0 s/mm^2^) were also acquired. Participants also underwent high-resolution T1-weighted magnetization prepared rapid gradient echo scanning (MPRAGE: echo time 3.06 ms; repetition time 2250 ms sequence, flip angle 9°, field of view=256 × 256 mm, acquisition matrix 256 × 256, voxel size 1 × 1 × 1 mm).

### DWI data processing and modelling

DWI data were converted from DICOM to NIfTI format using dcm2nii (www.nitrc.org/projects/dcm2nii). The images were skull-stripped, corrected for eddy currents and head motion using FSL (FSL 5.0.9 www.fmrib.ox.ac.uk/fsl). DWI data from both shells were registered to the first non-diffusion (*b* = 0 s/mm^2^) volume.

After pre-processing, diffusion tensors were fitted to the DWI data using DTIFIT in FSL for each shell (*b*=1200 s/mm^2^ and *b*=2400 s/mm^2^). Local fiber orientation distributions in each voxel were estimated by using the ball and stick model for multiple shells (Behrens et al., 2003; Jbabdi et al., 2012). For each voxel, we calculated two diffusion tensor imaging (DTI) measures, fractional anisotropy (FA) and mean diffusivity (MD) by using the *b*=1200 s/mm^2^ shell data (Fig. 2A). FA is a scalar ranging from 0 to 1 that quantifies the coherence of water diffusion, with 0 indicating low coherence and 1 indicating high coherence, while MD measures the average rate of water diffusion, with higher rates indicating fewer boundaries for water diffusion (Basser and Pierpaoli, 1996).

**Figure 2.**
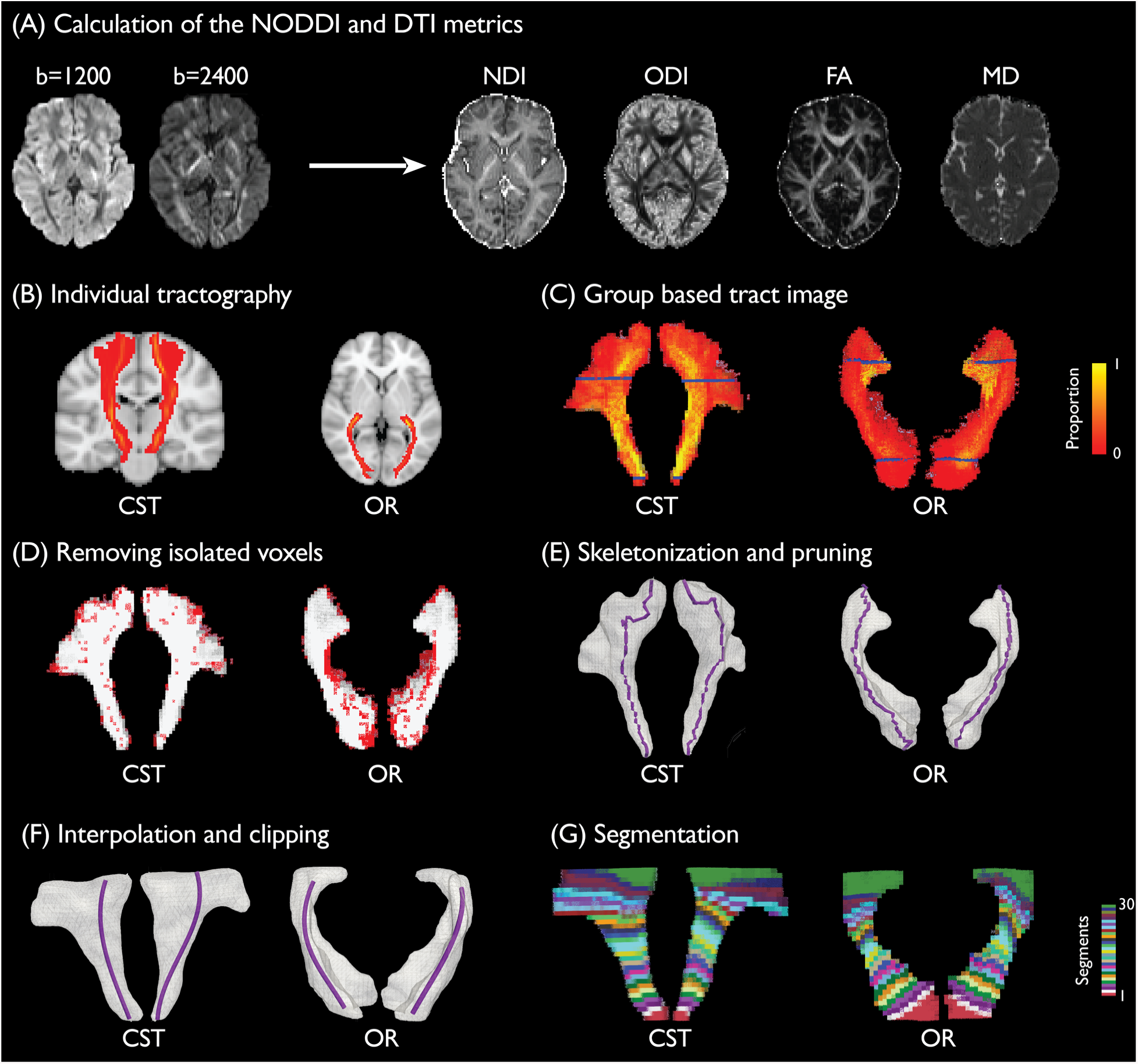
Analysis pipeline of along tract microstructural metrics. **(A)** After pre-processing, whole brain voxel-wise NODDI (NDI and ODI) and DTI (FA and MD) metrics were calculated from the two-shell DWI data. **(B)** Probabilistic tractography of CST and OR were calculated and thresholded in individual participant’s native space. **(C)** Individual tractography results of CST and OR were normalized to the MNI space and united to obtain group-based tract images. The voxel intensity in the group-based tract images denotes the proportion of overlapping across participants. **(D)** Isolated voxels (denoted as red) with no second order neighbors were removed from the group-based tract volumes. **(E)** Three-dimensional group-based tract volume image was skeletonized using thinning algorithms. Subsidiary branches of the skeleton were trimmed. **(F)** The main skeleton was smoothed with cubic spline interpolation. The central portion of the group-based tract volume (between the two blue planes shown in ***C***) and the skeleton were clipped for further processing. **(G)** The skeletons of the CST and OR were divided into 30 segments with equal lengths and 20% overlap between adjacent segments. Voxels in each group-based tract volume were assigned to the nearest segment based on Euclidean distance, resulting in 30 equidistant sub-volumes along the principal skeleton. Each sub-volume was transformed back to the individual’s native space to calculate microstructural metrics along tracts.

The same pre-processed DWI data was fitted to the Neurite Orientation Dispersion and Density Imaging (NODDI) model using the NODDI toolbox (www.nitrc.org/projects/noddi_toolbox). The NODDI model contains three tissue compartments: intra-cellular space, extra-cellular space and cerebrospinal fluid (CSF). The intra-cellular estimation uses the stick model to capture the restricted diffusion perpendicular to neurites and unhindered diffusion along them. The extra-cellular compartment models the hindered diffusion of water molecules by Gaussian anisotropic diffusion with parallel and perpendicular diffusivities. The CSF compartment is modelled as isotropic Gaussian diffusion to minimize the confounding effect of CSF contamination (Zhang et al., 2012a). From the model-derived intra-and extra-cellular compartments, we calculated two voxel-wise NODDI measures, Neurite Density Index (NDI) and Orientation Dispersion Index (ODI) (Fig. 2A). NDI represents the volume fraction of the intracellular compartment that contains the axons and dendrites, and ODI quantifies the angular variation in neurite orientation.

The high-resolution T1-weighted MPRAGE image was linearly co-registered to the native DWI space using mutual information with 6 degrees of freedom. The co-registered MPRAGE image was segmented and normalized to the Montreal Neurological Institute (MNI) standard template by linear and non-linear deformations using FSL. The forward and inverse deformation fields between the native DWI space and the MNI template space were used for subsequent tractography and along-tract analysis.

### Tractography

For each participant, we conducted probabilistic tractography to reconstruct bilateral corticospinal tracts (CST) and optic radiation (OR) in the individual’s native space using FMRIB’s Diffusion Toolbox (Behrens et al., 2007). For both tracts, we used multiple regions of interest (ROI) from the Jülich histological atlas (Bürgel et al., 2006) and John Hopkins University (JHU) DTI-based white matter atlases (Hua et al., 2008) to define seed masks, target masks, waypoints and exclusion masks.

Each tract was reconstructed by sampling 5000 streamlines per voxel with 0.5 mm step length, 0.2 curvature threshold, 0.1 fiber threshold and 3 mm minimum streamline length. The probabilistic tractography procedure generates an image in which the intensity of each voxel is the ratio of number of streamlines that pass through that voxel over the total number of streamlines generated in the seed voxels (Fig. 2B). We thresholded the probabilistic tractography outputs at 5 × 10^−6^ to discard false negatives (Rilling et al., 2008).

For the CST, we followed a previous method (Zhang et al., 2010) to use the ROIs from the JHU atlases. The tractography was seeded from the cerebral peduncle (CP) and extended through the posterior limb of interior capsule (PLIC) and superior corona radiata (SCR). The target mask was the intersection of the precentral gyrus with the individual grey matter images. The target mask was dilated with a 2mm disk-shaped kernel using the Image Processing Toolbox in Matlab to account for anatomical variability across individuals (Clatworthy et al., 2010). Exclusion masks included the contralateral hemisphere, anterior limb of the internal capsule, the retrolenticular part of the internal capsule, the pontine crossing tract, the inferior and superior cerebellar peduncle.

For the OR, the tractography was seeded from the lateral geniculate nucleus (LGN) in each hemisphere, and a central slice of OR defined in the Jülich histological atlas was used as a waypoint mask to avoid the fibers that transverse Meyer’s loop (Clatworthy et al., 2010). The target mask was the primary visual cortex, jointly defined by the Jülich histological atlas and the individual grey matter mask from MPRAGE segmentation. This target mask was further dilated with a 2mm disk-shaped kernel to account for the individual anatomical variability. Exclusion masks included the anterior boundary of the OR in the Jülich atlas and the contralateral hemisphere (Clatworthy et al., 2010).

After tractography, we generated a group-based representation of each tract in the MNI space across all participants (Fig. 2C, Fig. 3A and 4A). The individual thresholded tractography results were binarized following the transformation to the MNI space by applying the forward deformation field from MPRAGE normalization. A group-based tract image was calculated from the union of all participants’ normalized tracts.

**Figure 3.**
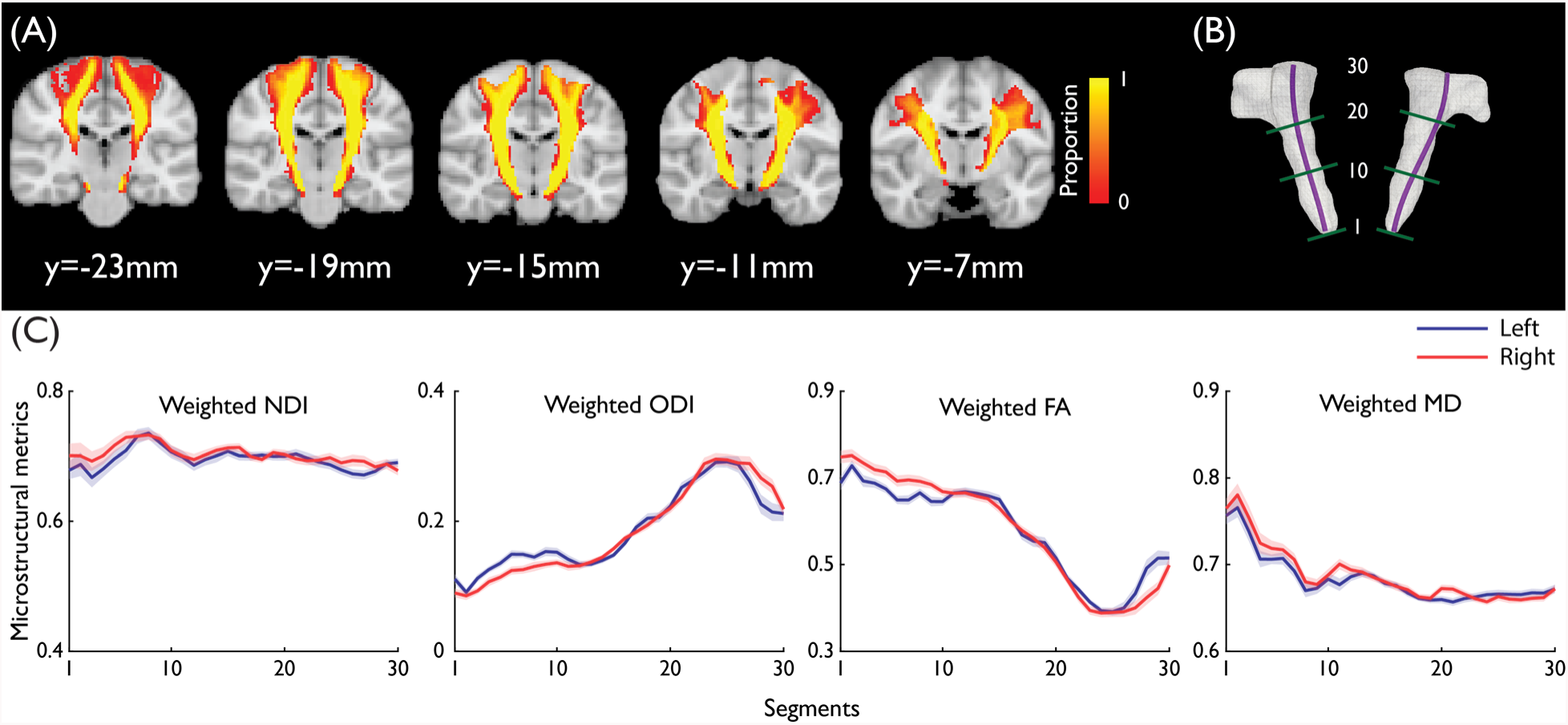
**(A)** Group-based image of the corticospinal tract (CST). The voxel intensity denotes the proportion of overlapping across participants. **(B)** Visualization of the CST with the skeleton and the approximate locations of segments 1, 10, 20 and 30. **(C)** The means of NDI, ODI, MD and FA profiles of the left and right CST across participants. Shaded areas represent 95% CI. The segments 1 to 30 were from Cerebral Peduncle to Superior Corona Radiata as in ***B***.

**Figure 4.**
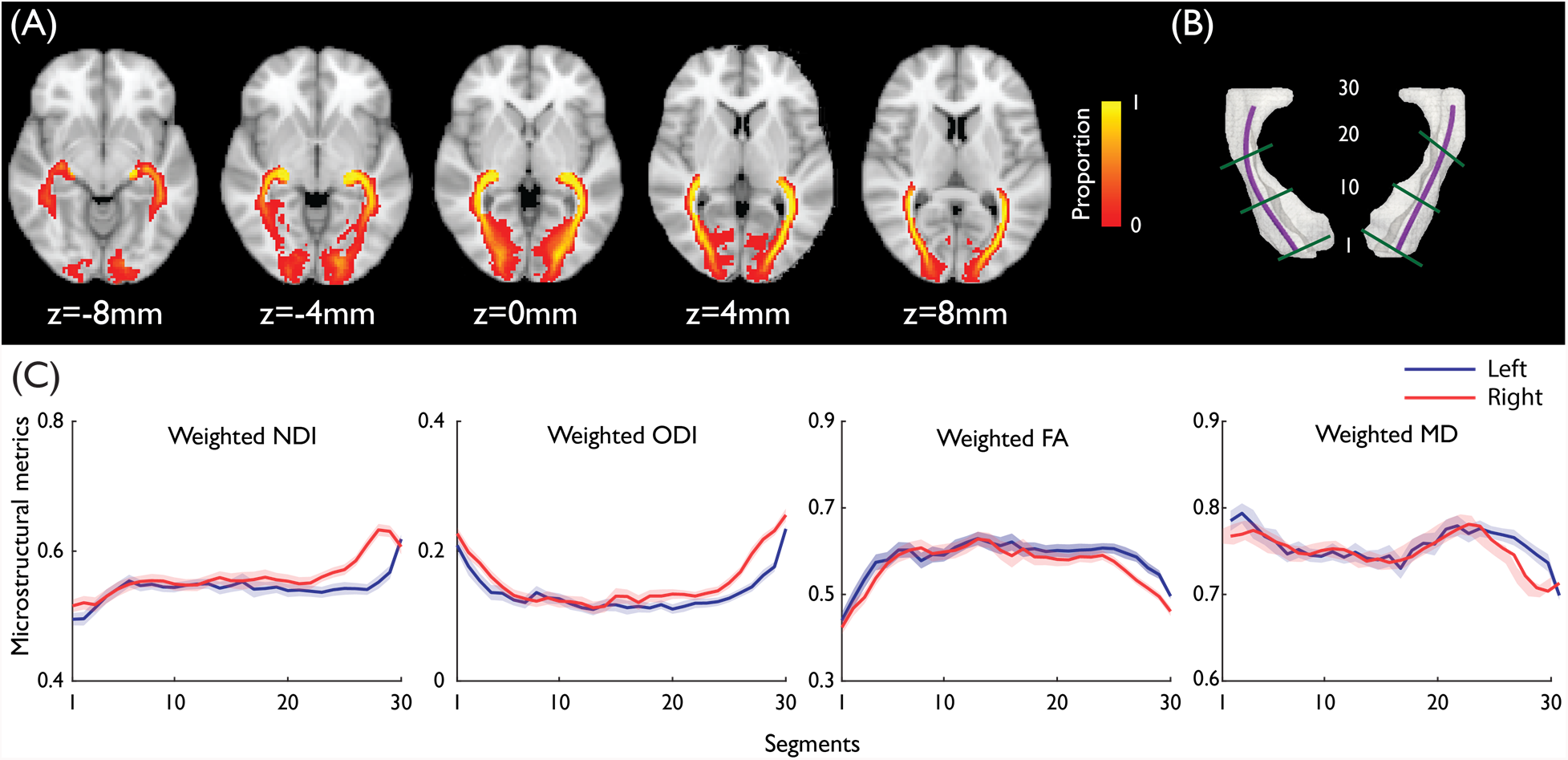
**(A)** Group-based image of the optic radiation (OR). The voxel intensity denotes the proportion of overlapping across participants. **(B)** Visualization of the OR with the skeleton and the approximate locations of segments 1, 10, 20 and 30. **(C)** The means of NDI, ODI, MD and FA profiles of the left and right OR across participants. Shaded areas represent 95% CI. The segments 1 to 30 were from V1 to posterior lateral geniculate nucleus as in ***B***.

### Tract skeletonization

For both CST and OR, we generated a skeletal profile of the group tract images using in-house scripts and the volume skeleton toolbox in Matlab (Fig. 2D, see also (Cornea et al., 2007)). First, the group tract image was binarized. The isolated voxels with no second order neighbors were removed. Holes and discontinuities in the tract image volume were removed by dilating the whole tract volume with a 2mm disk-shaped kernel. Second, we applied the thinning algorithm to estimate the skeleton of the cleaned tract images (Cornea et al., 2007). The thinning algorithm estimates the curve-skeleton of a three-dimensional object by iteratively removing simple voxels from the surface boundary. Simple voxels are defined as the ones whose removal would not change the topology of the volume. This operation was repeated until no simple voxels are remained. To find the main curvature of the tract, branches along the main trunk of the skeleton were removed. A cubic spline interpolation was fitted to the principal trunk of the skeleton to obtain a smoothed three-dimensional skeleton of the tract. Third, similar to previous approaches (Yeatman et al., 2012), we focused on the central portion of the tracts, where fiber bundle morphology is most consistent across individuals. Therefore, we clipped the group-based CST image and the skeleton to the portion between the cerebral peduncle and the superior corona radiata, and we clipped the group-based OR image and the skeleton to the portion closer to anterior regions of V1 and posterior regions of LGN.

### Microstructural measures along tracts

To quantify microstructural metrics and their changes along tracts, we divided the spline interpolation of each group-based tract skeleton into 30 segments with equal length and 20% overlap between adjacent segments, although our main results were not dependent on the exact number of segments. All the voxels in the group-based tract image were assigned to the corresponding segment based on the closest Euclidean distance. This procedure was repeated for all the four tracts (CST and OR in the left and right hemispheres), resulting in 30 equidistant sub-volumes along the principal skeleton of each group-based tract image (Fig. 2E). Because adjacent segments were overlapping, the labelled sub-volumes also had overlapping voxels. This interpolation ensured the along tract microstructural measures were less affected by segment boundaries (Yeatman et al., 2012).

All the sub-volumes were transformed back to the individual’s native space to reconstruct participant specific sub-volumes along tracts. For each participant, a voxel in a sub-volume was included in the calculation of individual microstructural metrics only if: (1) it was in the individual white matter mask from MPRAGE segmentation; (2) it had a FA value greater than 0.2 from DTI modelling; and (3) it had a suprathreshold probability of connection from the tractography results. The combination of group-level segments and individual-level constraints balances the consistency and variability across participants when making inferences along tracts.

For each participant, the four microstructural metrics (NDI, ODI and FA, MD) at each sub-volume of each tract were calculated as a weighted average of the metrics from all voxels within the sub-volume *M*:

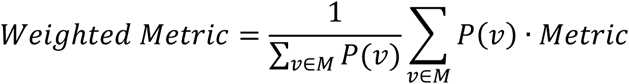

where the weight *P(v)* for voxel *v* is the normalized probability of connection from tractography, given by:

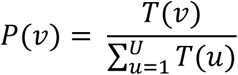

where *U* is the total number of the voxels in the brain accounted for tractography and *T(v)* is the probability of connection.

### Statistical analysis

To examine the relationships between RT and *T_er_*, we used both frequentist and Bayesian Pearson’s correlation test across participants (JASP team, 2018). We used repeated-measures ANOVA with two factors of track segments and hemispheres to examine the change of microstructural measures along tracts and across hemispheres. Greenhouse-Geisser correction was applied where Mauchly‘s Sphericity test indicated that the assumption of sphericity was violated.

For each sub-volume along a tract, we used general linear models to associate the weighted average of microstructural metrics to individual mean RT and *T_er_* from the behavioral data. Because of the age-related changes in DWI signal (Yeatman et al., 2014; Cox et al., 2016), we included age as a nuisance variable in all the models. To correct for multiple comparisons from the 30 overlapping segments along each tract, we used threshold-free cluster enhancement in FSL with 10,000 permutations to control family-wise error in each tract and microstructural metric (Smith and Nichols, 2009; Winkler et al., 2014).

### Software Accessibility

The algorithms for tract skeletonization, tract segmentation and other analyses used in the current study was open-source and freely available online (https://github.com/esinkarahan/ATA).

## Results

We examined whether individual-differences in brain microstructure would be associated with the reaction time of simple motor actions. Participants were required to perform a right-hand simple reaction time (SRT) task (Fig. 1A) to measure their mean RT (Fig. 1C). Next, we used the LBA model (Brown and Heathcote, 2008) to quantify the non-decision time *T_er_* during simple actions (Fig. 1B), a model-derived parameter to account for the latency of motor initiation and stimulus encoding (Lo and Wang, 2006; Cavanagh et al., 2011; Donkin et al., 2011). The two behavioral measures were then associated with microstructural metrics from NODDI and DTI models in the primary motor and visual pathways: the corticospinal tract (CST) and optical radiation (OR).

### Behavioral results

The mean RT across 46 participants was 363.02 ± 30.832 ms (S.D.) and the mean *T_er_* was 249.238 ± 48.227 ms (S.D.) (Fig. 1C). The LBA model provided an adequate fit to the observed RT distributions (Fig. 1D). We used frequentist and Bayesian Pearson’s correlation to examine the relations between behavioral measures. There was no significant correlation between mean RT and *T_er_* (*R* = 0.114, *p* = 0.45, 95% CI = [−0.182, 0.392], BF_10_ = 0.243), and no correlations between the behavioral measures and age (RT: *R* = 0.074, *p* = 0.624, 95% CI = [−0.221, 0.357], BF_10_ = 0.206; *T_er_*: *R* = 0.062, *p* = 0.683, 95% CI = [−0.233, 0.346], BF_10_ = 0.199).

### Tractography and along-tract microstructural metrics

We used a ROI-based probabilistic tractography approach to reconstruct four tracts (left CST, right CST, left OR and right OR) in the individual’s native space and obtained group-based tract images after normalizing individual tracts to the MNI space. From the group-based tract images (Fig. 3A), the inferior CST close to the brainstem showed high consistency across participants and the superior CST had large individual variability when it approaches the cortex. Similarly, there was a large tract variability where the OR approaches V1 (Fig. 4A).

We calculated the weighted NODDI (ODI, NDI) and weighted DTI (FA, MD) metrics from the 30 segments of each tract, equidistant along the tract’s principal skeleton (Fig. 3C and 4C). A repeated-measures ANOVA on microstructural metrics showed significant main effects of segments in CST (NDI: *F*(5.04, 226.65) = 20.82, *p* = 3.44E-17; ODI: *F*(4.18,188.28) = 550.94, *p*= 3.01E-104; FA: *F*(5.47,246.29) = 730.35, *p* = 1.408E-149; MD: *F*(5.14,231.08) = 168.80, *p* = 5.48E-76, Greenhouse-Geisser corrected) and OR (NDI: *F*(8.01,360.37) = 64.13, *p* = 9.99 E-65; ODI: *F*(8.08,363.42) = 126.63, *p* = 1.1E-100; FA: *F*(8.72,392.55) = 71.62, *p*=1.01E-245; MD: *F*(11.65,524.08) = 32.38, *p* = 9.75E-132).

In the CST, there was significant hemispheric difference in NDI (*F*(1,45) = 18.35, *p* = 0.000095) and MD (*F*(1,45) = 17, *p* = 0.00016) and the hemispheric difference in ODI and FA did not reach significance (ODI: *F*(1,45) = 3.81, *p* = 0.057; FA: *F*(1,45) = 1.85, *p* = 0.18). In the OR, there was significant hemispheric difference in all metrics (NDI: *F*(1,45) = 61.18, *p*=6.27E-10; ODI: *F*(1,45) = 34.89, *p*=4.29E-7; FA: *F*(1,45)=15.22, *p*=.00032; MD: *F*(1,45)=6.7, *p*=0.013).

These main effects were qualified by significant interactions between segment locations and hemispheres in CST (NDI: *F*(6.04,271.79) = 3.569, *p* = 0.002; ODI: *F*(6.43,289.33) = 20.13, *p* = 1.15E-20; FA: *F*(8.78,395.14) = 23.41, *p* = 1.98E-31; MD: *F*(9.15,411.59) = 4.4, *p* = 0.000014) and OR (NDI: *F*(9.55,429.75) = 17.862, *p* = 4.134E-75; ODI: *F*(10.39,467.53) = 9.72, *p* = 2.995E-15; FA: *F*(9.43,424.34) = 3.8, *p* = 0.000095; MD: *F*(11.43,514.51) = 4.76, *p* = 3.59E-7). Therefore, there were substantial variations in microstructural metrics along CST and OR.

We examined the correlations between the microstructural metrics along tract segments (Fig. 5). Threshold-free cluster enhancement (TFCE) with 10,000 permutations was used to correct multiple comparisons for the number of segments along tracts. The NODDI measure of NDI was positively correlated with ODI in superior segments of the CST and anterior segments of OR, and ODI was negatively correlated with FA and MD in most of the segments, consistent with previous results (Zhang et al., 2012a). FA and MD showed both positive and negative correlations in some segments, which may be due to changes in local tract geometry along the tracts.

**Figure 5.**
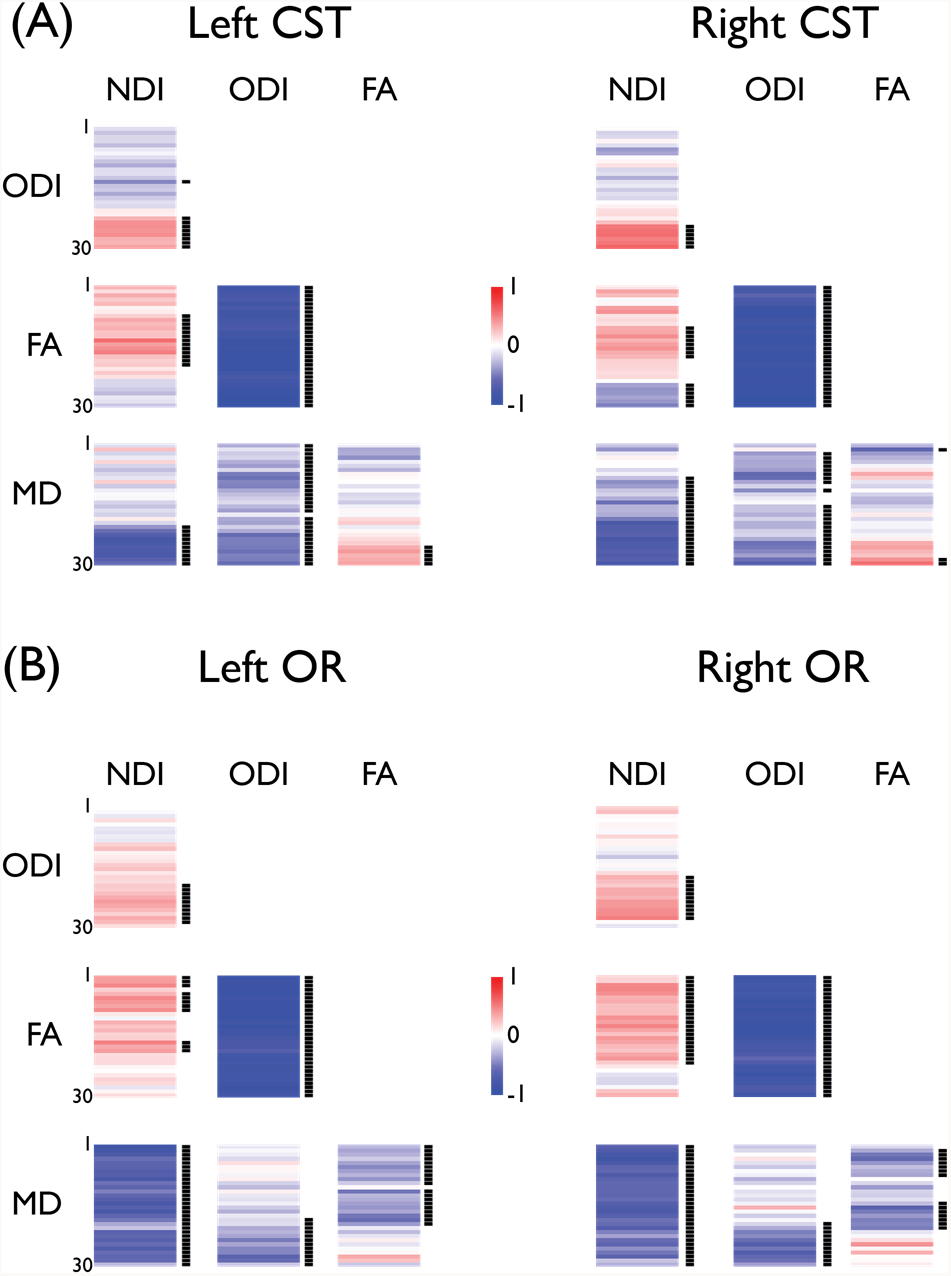
Correlation between NODDI and DTI metrics along the CST **(A)** and OR **(B)** segments across participants. As in Figs. 3B and 4B, the anatomical locations of the segments were from inferior to superior for CST, and posterior to anterior for OR. Positive and negative Pearson’s correlation coefficients are color-coded separately. Bars at right of each graph indicate segments with significant correlations (*p*<0.05, 10,000 permutations) after threshold-free cluster enhancement correction for the number of segments.

### Correlating response speed measures with tract microstructure

We used general linear models to examine the associations between response speed measures (SRT and *T_er_*) and microstructure metrics in all tract segments, including age as a nuisance variable. Threshold-free cluster enhancement (TFCE) with 10,000 permutations was used to correct multiple comparisons for the number of segments along tracts.

For the CST (Fig. 6A), faster *T_er_* was associated with higher NDI values in the superior segments of both left (segments 18-30, *p*<0.05, TFCE corrected) and right tracts (segments 15-28, *p*<0.05, TFCE corrected). The segments with significant correlations comprised the posterior limb of interior capsule and the superior corona radiata that connect to the precentral gyrus (Fig. 7A). No other microstructural metrics of the CST had significant correlation with the *T_er_* (across all segments of the left CST: *p*>0.331; right CST: *p*>0.116).

**Figure 6.**
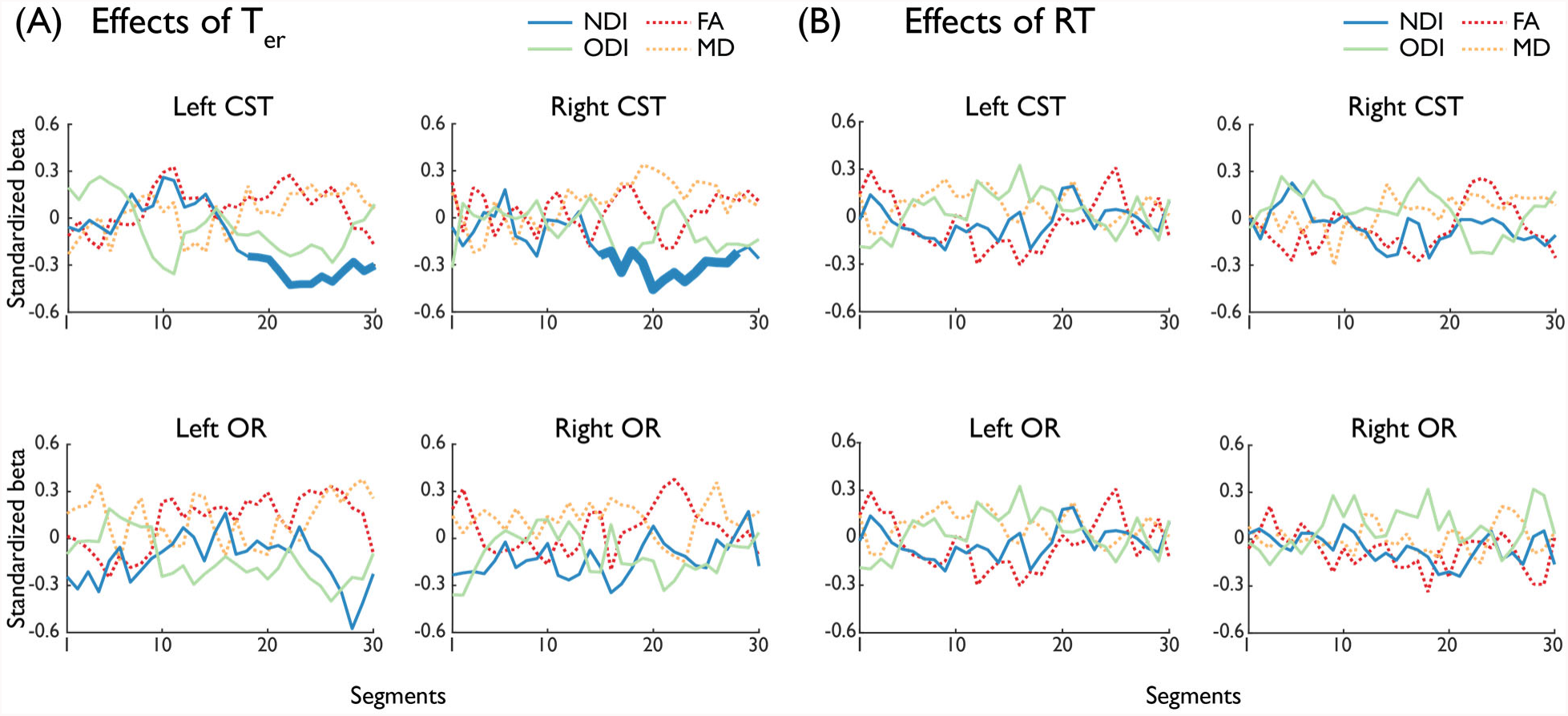
Standardized linear regression coefficients between microstructural metrics and *T_er_* (**A**) and mean RT (**B**). the behavioral parameters. The segments with significant associations after threshold-free cluster enhancement correction (*p*<0.05, 10,000 permutations) are presented with increased line thickness.

**Figure 7.**
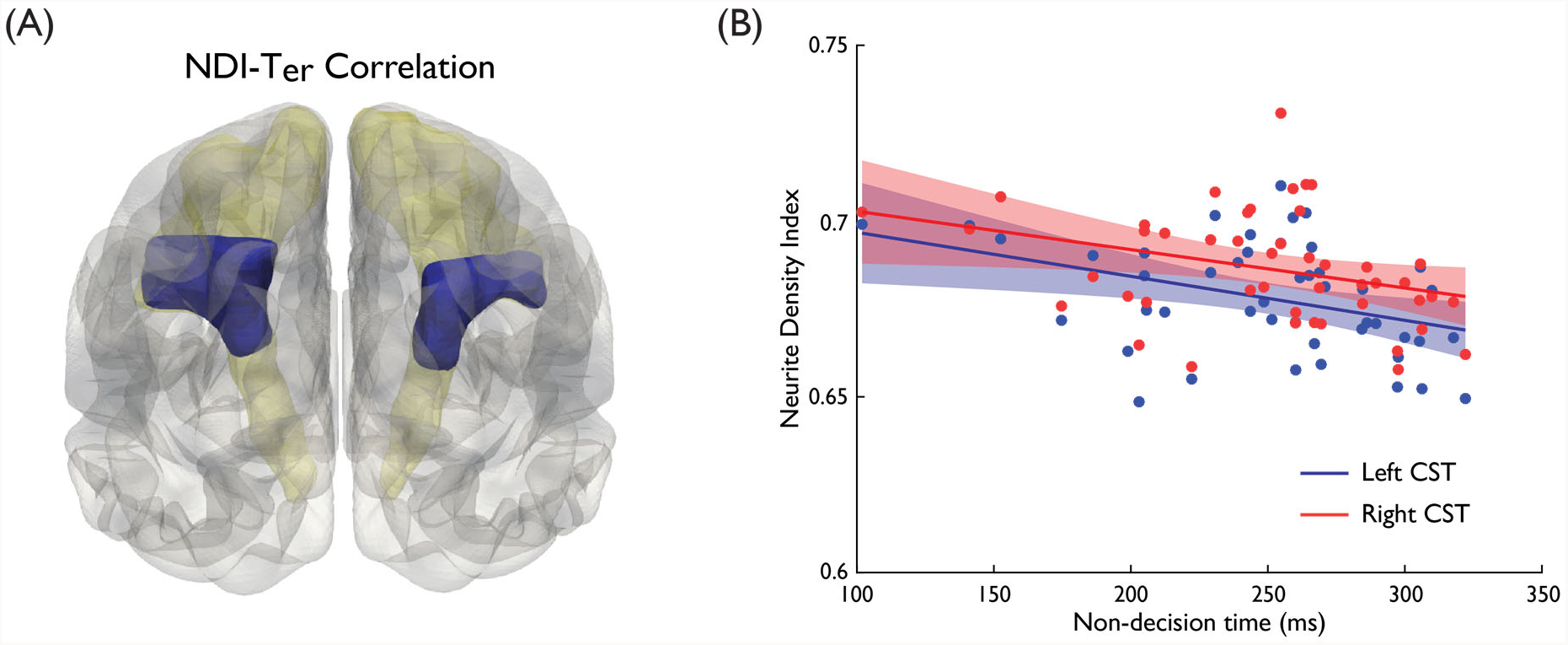
(**A**) CST segments that exhibited a significant negative NDI-*T_er_* correlation are highlighted (in blue) over the group-based tract volume. (**B**) Correlation between the *T_er_* and the NDI from the entire left (blue) and right (red) CST. Each data point represents an individual participant. Shaded areas represent the 95% confidence intervals.

For the OR (Fig. 6A), there was no significant correlation between *T_er_* and any microstructural metrics after correction of multiple comparison (across all segments of the left OR: *p*>0.075; right OR: *p*>0.24). At an uncorrected threshold of *p*<0.01, *T_er_* was positively correlated with the FA in segment 27 of the left OR.

There was no significant correlation between mean RT and microstructural metrics in CST (across all segments of the left CST: *p*>0.425; right CST: *p*>0.31) or OR (across all segments of the left OR: *p*>0.108; right OR: *p*>0.479) (Fig 6B).

To examine further in a *post-hoc* exploratory analysis whether the significant *T_er_*-NDI associations in CST extended to the entire tract, we calculated the NDI of the whole CST and its partial correlations with the *T_er_*, controlling for the effect of age. There were significant negative correlations between the *T_er_* and the NDI values of bilateral CST (Fig. 7B; left CST: *R* = −0.390, *p*= 0.008, 95% CI = [−0.582, −0.157], BF_10_ = 5.118; right CST: *R* = −0.341, *p* = 0.022, 95% CI = [−0.544, −0.101], BF_10_ = 2.097).

## Discussion

By combining cognitive and biophysical models with novel along-tract analyses in healthy adults, we found that the neurite density index (NDI) of superior segments of bilateral CST negatively correlated with the non-decision time (*T_er_*), a model-derived component of RT that represents the duration of non-decision processes in simple actions (Lo and Wang, 2006; Cavanagh et al., 2011; Donkin et al., 2011). At an uncorrected threshold, *T_er_* positively correlated with the FA in an anterior segment of the left OR. Together, our findings provide new insights into how microstructural properties in white matter pathways contributed to inter-individual differences in response speed in young adults.

The compartmental model of NODDI quantifies intracellular neurite density separately from orientation dispersion (Zhang et al., 2012a), which are otherwise indistinguishable in standard DTI measures (e.g., FA). The specific tissue property measures offered by NODDI are sensitive to microstructural changes in previous studies of brain development (Kunz et al., 2014; Chang et al., 2015; Kodiweera et al., 2016; Genc et al., 2018), psychosis (Rae et al., 2017) and neurodegeneration (Colgan et al., 2016; Slattery et al., 2017). We demonstrated that faster non-decision time was associated with higher NODDI-derived neurite density index in the superior segments of bilateral CST, including a part of the superior corona radiata and its connections to the precentral gyrus. This region had a large dispersion of fiber orientation, as indicated by the large ODI values (segments 14-30, Fig. 3C) and positive correlation between ODI and NDI metrics (segments 23-30 in left CST and segments 25-30 in right CST, Fig. 5A). The significant correlation observed in this region may relate to somatotopic organization of the CST (Hong et al., 2010; Seo et al., 2012) and distal (finger) movements used in the task, which needs to be confirmed in future studies combining functional localizers and tractography (Dalamagkas et al., 2018).

In humans, neurite density estimates from diffusion MRI correlate strongly with cortical myelin content (Fukutomi et al., 2018) and are sensitive to demyelination disorders such as multiple sclerosis (Grussu et al., 2017; Schneider, 2017). In rodents, neurite density estimates have a good agreement with the intensity of myelin stain under light microscopy (Jespersen et al., 2010). Therefore, although the NDI does not distinguish between myelin and axonal volume fraction (Jelescu et al., 2015; Stikov et al., 2015), the negative *T_er_*-NDI correlation in the current study supports the hypothesis that participants with faster motor speed have thicker myelin sheaths or axonal diameter in the CST, which in turn results in faster nerve conduction velocity (Waxman, 1980; Fields, 2008; Seidl, 2014; Buskey et al., 2017).

In the left OR, the positive correlation between FA and *T_er_* did not survive the correction for multiple comparisons. Nevertheless, previous studies using a voxel-wise approach reported similar associations between FA and choice RT in OR (Tuch et al., 2005). The SRT task in the current study may not be sensitive to the inter-individual variability in the visual processing pathway, as suggested by the lack of change of early visual event-related potential in the SRT task compared with a choice RT task (Mangun and Hillyard, 1991).

Our study provides new methods for studying brain, behavior and cognition relationships. First, we used microstructural measures weighted by connection probability and along tract analysis based on volumetric skeletonization and segmentation. Similar to previous studies (Yeatman et al., 2012), we showed significant variations in all DTI and NODDI measures along CST and OR. These variations may be due to changes in microscopic tissue properties (Murray and Coulter, 1981), local tract geometry, or neighboring environments such as partial volumes and crossing fibers. More importantly, correlations with behavioral measures were observed only in a portion along the tracts, confirming the needs to take into account microstructural variations along tracts (Yeatman et al., 2014). Several methods have previously been proposed for characterizing microstructural metrics along tracts, including medial axis representations (Yushkevich et al., 2008; Cheng et al., 2012), b-spline resampling of streamlines (Colby et al., 2012), arc length matching to prototype fibers (O’Donnell et al., 2009) and centroid fibers based on the minimum (Wang et al., 2016) or mean (Yeatman et al., 2012) distance of fibers within a tract.

The current study co-registered individual participant’s tractography results to a template space and generated group-level tract probability maps (Hua et al., 2008), from which the tract skeleton and equidistant segments were calculated for subsequent along tract analysis. In both CST and OR, voxels close to the tract skeleton had high tract probabilities in the group maps (Fig. 3A and 4A), suggesting a good agreement across participants. This approach allows reduction of the noise from individual tractography and obtain representative volumetric tract profiles. Nevertheless, normalization errors and variances of tract geometries across participants may hamper the precision of inference at the individual level. To address this issue, we transformed segments along the tract skeleton back to the native space and accounted for individual heterogeneity using masks from individual tissue probability and tractography. Furthermore, in contrast to simply averaging across voxels, we capitalized on the connection distributions from probabilistic tractography to downweight the contributions from voxels with high noise or low certainty in the calculation of microstructural measures (Olvet et al., 2016). As a result, our method combined group-level along tract profiling as well as individual-level fiber tracking.

Our automated analysis pipeline can be applied to other tracts and suit to larger datasets. We note that, like other methods for along tract analysis, further tests are needed to ensure the robustness in smaller tracts with more substantial variability than demonstrated here, which beyond the scope of the current study. To facilitate future research, we have made our analysis scripts open source and freely available.

Second, our study highlighted the benefits of computational modelling of behavioral data (Forstmann et al., 2016). The LBA model decomposed RT distributions into the duration of an evidence accumulation process (Gold and Shadlen, 2007) and non-decision time *T_er_*. Individual differences in *T_er_* are likely subject to influences at multiple processing levels. Along CST and OR, we found that microstructural metrics correlated with the *T_er_* but not mean RT. Considering the predominate roles of these tracts in transmitting motor and visual information, our results provide anatomical evidence to support the common assumption that the *T_er_* includes motor execution and visual encoding latencies. The *T_er_* is a unitary estimate and hence cannot distinguish between visual and motor latencies. It is possible to combine our approach with other imaging modalities to dissociate the *T_er_* into subcomponents that occur before and after decisions (White et al., 2014; Nunez et al., 2018).

There are several limitations to this study. Our experimental samples included only female participants with a narrow age range. Both gender (Maccoby, 1991; Dykiert et al., 2012) and age (Wilkinson and Allison, 1989) have been shown to affect RT (Yang et al., 2015). A cross-sectional study of over 3,000 participants showed that gender and age also affect NODDI and DTI measures in multiple white matter tracts (Cox et al., 2016), in line with other smaller-sample studies (Good et al., 2001; Toosy et al., 2004; Lebel et al., 2012; Bede et al., 2014; Kodiweera et al., 2016). Further research could extend our results to heterogeneous or genetically-informed samples to examine the source of inter-individual variability, because both white-matter microstructure and RT are affected by genetic and environmental factors (Martin, 2004; Chiang et al., 2011; Rüber et al., 2015).

We focused on two *a priori* tracts of primary motor and visual pathways: CST and OR, because the latencies of motor initiation and stimulus encoding are commonly assumed to comprise the *T_er_*. This does not rule out the possibility that RT in other behavioral tasks associate with microstructural measures in different fiber tracts. Indeed, the RT in a visual oddball task correlated with the MD in tracts connecting the visual cortex with frontal and temporal cortices (Konrad et al., 2009), and the RT in a selective attention task correlated with the FA in the superior and inferior longitudinal fasciculus (Mayer and Vuong, 2014). The current study only estimated the *T_er_* and mean RT from a single SRT task, and hence cannot infer the extent to which the microstructural correlates of the response speed may vary as a function of different tasks. Furthermore, other imaging sequences and models are now available to obtain histologically validated microstructural measures beyond NODDI, such as axon diameters and *g*-ratios (Deoni et al., 2008; Alexander et al., 2010; Thiessen et al., 2013). An important future direction is to have a mechanistical understanding of how quantitative tissue microstructure may selectively influence behavioral performance across cognitive domains.

In conclusion, inter-individual differences in non-decision time during simple actions were reflected in the extent of neurite density in white matter tracts responsible for motor information transmission. These findings helped to validate the functional origin of non-decision time assumed in current models of decision making and action selection. Our results further raised an intriguing possibility that tissue microstructure in key fiber tracts relate to response speed in basic cognitive processes.

## Acknowledgements

This study was supported by a European Research Council Starting grant (716321). AGC’s PhD studentship was funded by the Cardiff University Neuroscience and Mental Health Research Institute. KSG and ADL were supported by the Wellcome Trust Strategic Award (104943/Z/14/Z).

